# Selection on morphological traits and fluctuating asymmetry by a fungal parasite in the yellow dung fly

**DOI:** 10.1101/136325

**Authors:** Wolf U. Blanckenhorn

## Abstract

A Preprint reviewed and recommended by **Peer Community Evolutionary Biology**: http://dx.doi.org/10.24072/pci.evolbiol.100027

Evidence for selective disadvantages of large body size remains scarce in general. Previous phenomenological studies of the yellow dung fly *Scathophaga stercoraria* have demonstrated strong positive sexual and fecundity selection on male and female size. Nevertheless, the body size of flies from a Swiss study population has declined by almost 10% from 1993 to 2009. Given substantial heritability of body size, this negative evolutionary response of an evidently positively selected trait suggests important selective factors being missed (e.g. size-selective predation or parasitism). A periodic epidemic outbreak of the fungus *Entomophthora scatophagae* allowed assessment of selection exerted by this parasite fatal to adult flies. Fungal infection varied over the season from ca. 50% in the cooler and more humid spring and autumn to almost 0% in summer. The probability of dying from fungal infection increased with adult body size. All infected females died before laying eggs, so there was no fungus impact on female fecundity beyond its impact on mortality. Large males showed the typical mating advantage in the field, but this pattern of positive sexual selection was nullified by fungal infection. Mean fluctuating asymmetry of paired appendages (legs, wings) did not affect the viability, fecundity or mating success of yellow dung flies in the field. This study demonstrates rare parasite-mediated disadvantages of large adult body size in the field. Reduced ability to combat parasites such as *Entomophthora* may be an immunity cost of large size in dung flies, although the hypothesized trade-off between fluctuating asymmetry, a presumed indicator of developmental instability and environmental stress, and immunocompetence was not found here.

## Introduction

Systematic quantification of selection has become one of the hallmarks of modernbiological research soas to acquire athorough understanding of the process of natural selection and its evolutionary consequences. Standardized measures of selection (Arnold & Wade, 1984a, b; Lande & Arnold, 1983; Brodie et al., 1995) have been available for some time and applied to many species and situations to foster multiple comparative (meta-)analyses, which greatly enhanced our understanding of the action of natural selection in the wild (e.g. Endler, 1986; Kingsolver et al., 2001; Kingsolver & Pfennig, 2004; Cox & Calsbeek, 2009). Correlative phenomenological investigations of selection also are the field method of choice to understand the evolution and population biology of single species or populations in an integrative way, and to test hypotheses about the evolution of particular traits and patterns (e.g. sexual size dimorphism: Blanckenhorn, 2007).

A common pattern is that in many species of animals fecundity selection favors large females and sexual selection large males (Roff, 1992; Andersson, 1994; Kingsolver & Pfennig, 2004). Large body size also confers a number viability benefits (e.g. a longer life), so the selective forces favoring small body size, presumably primarily related to juvenile viability costs of becoming large and/or costs of maintaining a large adult size, are often cryptic (Blanckenhorn, 2000). The widespread yellow dung fly *Scathophaga stercoraria* (Diptera: Scathophagidae) is a classic model species for studies of natural, particularly sexual selection (Parker, 1979; Borgia, 1982; SigurjÓnsdÓttir & Snorrason, 1995) and a typical case in point. Field studies have established temporally variable but on average very strong mating advantages of large males, as well as fecundity advantages of large females (Jann et al., 2000; Blanckenhorn et al., 2003; cf. Honek, 1993). Strong sexual selection on male body size likely is the main driver of the untypical male-biased sexual size dimorphism in this species (Fairbairn, 1997; Kraushaar & Blanckenhorn, 2002; Blanckenhorn, 2007, 2009). Nevertheless, the body size of flies from our field population in Switzerland has declined by almost 10% over a 15-year period from 1993 - 2009 (Blanckenhorn, 2015). Given generally substantial heritability of body size also in this species (Mousseau & Roff, 1987; Blanckenhorn, 2000), this negative evolutionary response of a trait that is measurably strongly positively selected suggests that we are missing important selective factors or episodes shaping the body size of yellow dung flies (Merilä et al., 2001; Blanckenhorn, 2015; Gotanda et al., 2015). In general, and also for the yellow dung fly, evidence for selective disadvantages of large body size remains scarce(Blanckenhorn, 2000, 2007).

One aspect not well studied in yellow dung flies is size-dependent survival in nature. This is generally the case for small-bodied invertebrates, for which longitudinal field studies are essentially impossible because individuals cannot be easily marked and followed in nature, as is the case for larger vertebrates (Merilä & Hendry, 2014, Schilthuizen & Kellermann, 2014; Stoks et al., 2014; Blanckenhorn, 2015). At the same time, laboratory longevity estimates (e.g. Blanckenhorn, 1997; Reim et al., 2006; Blanckenhorn et al., 2007) generally do not well reflect field mortality. Multiple field estimates of larval survivorship at various conditions exist suggesting some counter-selection against large body size via the necessarily longer development time (summarized in Blanckenhorn, 2007). However, sex-and size-specific adult survivorship in the field was so far estimated only indirectly by Burkhard et al. (2002) using age-grading by wing injuries. The present study assesses natural selection on morphological traits by the fatal fungal parasite *Entomophthora spp*.

The parasitic fungus *Entomophthora scatophagae* regularly infects yellow dung fly adults in Europe and North America (Hammer, 1941; Steinkraus & Kramer, 1988; Maitland, 1994; Steenberg et al., 2001). Primarily at humid conditions infections can be epidemic (pers. obs.), in which case infected dead flies can be found prominently exposed near cow pastures on flowers, long grass or fences in a characteristic posture presumably effectively disseminating fungal spores and/or attracting other flies (Maitland, 1994; Møller, 1993). Spore transmission likely also occurs via physical contact, e.g. during copulation (Møller, 1993). The fungus is fierce and effective at infecting and killing adult flies within hours or days and can be manipulated, such that related, often species-specific such fungi are being employed for biological control of some insect pests (e.g. Steenberg et al., 2001; Nielsen & Hayek, 2006). This study took advantage of a fungus epidemic at our field population near Zürich, Switzerland, in 2002.

I here assess viability, fecundity and sexual selection on morphology and fluctuating asymmetry. Morphological traits reflecting body size are often assessed in selection studies (Kingsolver et al., 2001; Kingsolver & Pfennig, 2004). Body size is one of the most important quantitative traits of an organism, as it strongly affects most physiological and fitness traits (Calder, 1984; Schmidt-Nielsen, 1984; Roff, 1992) and exhibits several prominent evolutionary patterns in many organisms (Rensch, 1950; Fairbairn 1997; Blanckenhorn, 2000; Blanckenhorn & Demont, 2004; Kingsolver & Pfennig, 2004). Depending on the taxon, diverse traits are typically used as surrogates of body size, which are usually highly correlated (i.e. integrated) within individuals due to pleiotropy, epistasis or gene linkage. Nevertheless, for functional reasons selection on various body parts may differ (e.g. Preziosi & Fairbairn, 2000; Fairbairn, 2005), producing responses in correlated traits and thus generally prompting a multivariate approach (Lande & Arnold, 1983). I focused on paired appendages (legs, wings), so I could also assess fluctuating asymmetry (FA; Palmer & Strobeck, 1986). Small and random deviations from the a priori perfect symmetry in bilaterally symmetric organisms, i.e. FA, are presumed to reflect heritable developmental instability, such that individuals with good genes and/or living in good conditions can produce more symmetric bodies in the face of environmental stress, ultimately augmenting their fitness. Symmetric individuals consequently should have greater survival prospects (viability selection), and should be more successful at acquiring mates (sexual selection: Møller & Swaddle, 1997). However, especially the latter notion, and the evidence, remain controversial(Møller & Thornhill, 1997; Palmer, 2000; Polak, 2003; Klingenberg, 2003; Van Dongen, 2006; Knierim et al., 2007). In yellow dung flies, Liggett et al. (1993) and Swaddle (1997) found a negative relationship between FA and mating and foraging success, respectively, while Floate & Coughlin (2010) found no evidence for FA being a useful biomarker of environmental stress exerted by toxic livestock medications (ivermectin). Our previous studies of this species found that FA is not heritable (Blanckenhorn & Hosken, 2003), that it does not increase with inbreeding (or homozygosity: Hosken et al., 2000), and that FA does not affect male mating success in the field, while nevertheless being negatively related to energy reserves (Blanckenhorn et al., 2003). Of key relevance here is the postulated link between FA and immunocompetence, predicting that more symmetric, but likely also larger individuals are better at fending off internal parasites such as *Entomophthora* (see e.g. Rantala et al., 2000, 2004, 2007 and Yourth et al., 2002, for various insect species), which was tested here.

## Material and Methods

### Study species

Yellow dung flies occur throughout the northern hemisphere and are particularly common around cow pastures in central Europe. The species prefers cooler climates. In lowland central Europe, each year has a spring (March – June) and an autumn season (September – November), while during the hot midsummer (July and August) the flies largely disappear from the pastures due to their heat sensitivity (Blanckenhorn, 2009).

Adult *S. stercoraria* are sit-and-wait predators of small flying insects, from which they extract protein to produce sperm and eggs (Foster, 1967). Females spend most of their time foraging for nectar and prey in the vegetation surrounding pastures. About once a week they visit fresh cattle (or other) dung to deposit a clutch of eggs. Larvae feed on and develop in the dung. Multiple males typically wait at the dung pat to mate with incoming females. Copulation usually takes place in the surrounding grass or on the dung pat; during the ensuing oviposition the male guards the female against other competitors (Parker, 1970). Competition among males for females is very strong as the operational sex ratio is highly male biased (Jann et al., 2000). Larvae face unpredictable spatio-temporal variation in local temperatures, dung (i.e. food) quality and quantity, intra-and inter-specific competition, and dung drying, all factors that ultimately largely determine their phenotypic adult body size. Towards the end of the season the flies have to reach the overwintering pupal stage before the first winter frost (Blanckenhorn, 2009).

### Fly sampling

I sampled our population in Fehraltorf near Zurich (N47°52’, E8°44’) roughly once a month between April and November 2002 (8 seasonal samples; total of *N*= 541 flies). Each time I sampled one randomly selected but otherwise representative fresh dung pat, collecting all single and paired flies on and ca. 20 cm around the pat to bring them alive to the laboratory. As virtually no unpaired females occur at a pat in the field because competition for mates is so intense, the number of pairs corresponds to the number of females present, and the proportion of paired males corresponds to the operational sex ratio (females / males).

### Laboratory procedures

Although dead flies infected with the fungus were occasionally found on and around the pasture, these were too rare and haphazard to be sampled systematically. Instead, all collected flies were kept alive in the laboratory in single 100 ml bottles with sugar and water for up to two weeks. Infected flies would develop the fungus within few days and eventually die; non-infected, recovered or resistant flies would not (acknowledging some random background mortality). Females received dung to oviposit one clutch of eggs, which was counted, which yields a good estimate of body size dependent clutch size (Honek, 1993; ca. 30 - 90 eggs: Jann et al., 2000; Blanckenhorn, 2007).

In the end, work study students measured left and right wing length as well as fore, mid and hind tibia length of each fly. Mean values for these paired traits were subsequently calculated, as well as signed FA as (L - R), unsigned FA as (|L - R|) (both in mm) and unsigned, size-corrected FA as (|L - R|) / mean(L, R) in %, as suggested by Palmer & Strobeck (1986). Paired traits were measured twice blindly by the same person to estimate measurement error relative to fluctuating asymmetry (Palmer & Strobeck, 1986), and to calculate the repeatability of all trait measurements (Becker, 1992). All measurements were taken with a binocular microscope at 16x magnification.

### Statistical analysis

For each monthly sample, and overall, I calculated standardized viability selection differentials(=gradients) for both sexes (binary variable: dead/alive = infected/uninfected), sexual selection differentials for males (binary: mated/unmated), and fecundity selection gradients for females based on their clutch size, using standard methods (Lande & Arnold, 1983; Arnold & Wade, 1984a,b; Brodie *et al.*, 1995). It turns out that almost all females that developed the fungus and eventually died in the laboratory did not lay any eggs, so fecundity selection coefficients only refer to healthy (uninfected) females. Selection coefficients were calculated for each trait (4 morphological and 4 asymmetry traits) separately for females and males, and additionally for the first principal component (PC) of all (mean) appendages signifying a compound index of body size, as such compound indices are often preferred by some researchers as proxies of overall body size even though they are redundant in this context.

Because the sizes of all appendages were highly positively correlated, and because FA and size are mathematically related (see formulae above), we calculated only univariate linear *(βuni)* and corresponding non-linear *(γ_uni_)* selection coefficients. To do so, for each seasonal sample we produced standardized *z*-scores for trait *x* by subtracting the sample mean from each value and dividing by the standard deviation: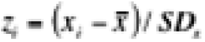. Relative survival or male pairing success was computed as absolute survival or pairing success (1 or 0) divided by the sample proportion of survived flies or mated males, respectively (Brodie & Janzen, 1996), and relative clutch size as absolute clutch size divided by mean clutch size. We used the univariate model of relative fitness on standardized body size 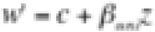 to estimate the linear selection intensities, and the corresponding quadratic model 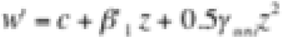 to estimate univariate non-linear selection intensities, 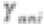. These linear coefficients (gradients) reflect the combined effects of direct and indirect selection on body size (Endler, 1986). The difference of the regression coefficients from a slope of zero (the null hypothesis of no selection) was tested. For estimation of the coefficients least-squares regression was applied, but for tests of significance logistic regression was used in case our measures of success were binary (viability and mating success: Brodie *et al.*, 1995; cf. Janzen & Stern, 1998). A full model with sex and season (including interactions) was performed to test for variation in selection across thesemain factors.

## Results

Fungus prevalence varied over the season and between the sexes. Infections (as high as 50%) were most common during the cooler and more humid periods at the beginning (spring) and the end of the season (autumn), whereas they were rare during the hotter summer (nearly 0%); females were more affected than males (significant sex by season interaction:***X*^2^** = 29.80; *P* < 0.001;Fig. 1). The probability of dying by fungal infection was unaffected by mean FA (***X*^2^** = 1.99; *P* = 0.159)but increased with fly body size (***X*^2^** = 12.56; *P* < 0.001; Table 1), an effect that however varied among seasonal samples (***X*^2^** =12.55; *P* < 0.001) but not between the sexes (***X*^2^** = 0.27; *P* = 0.602).

**Figure 1:**
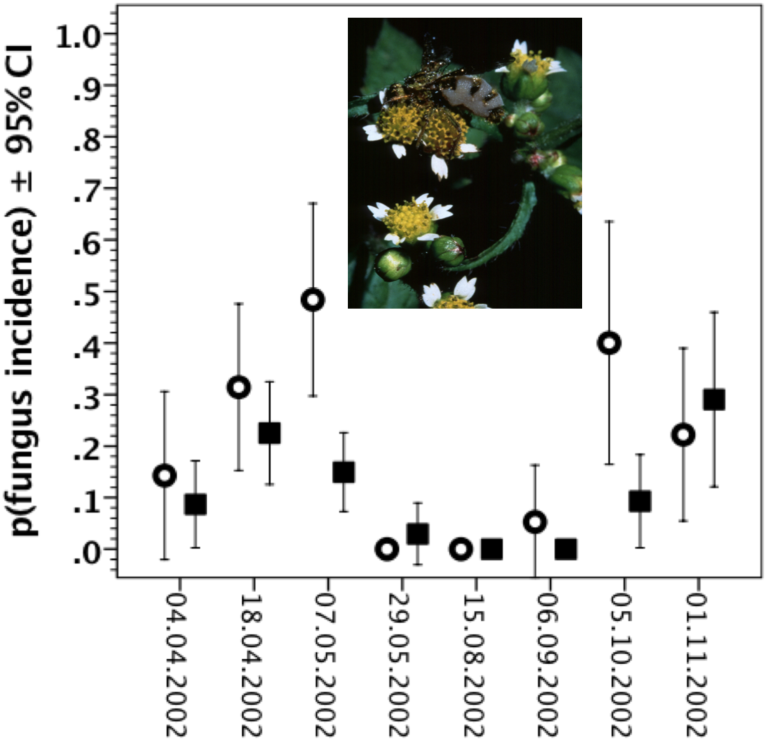
Proportion of male (filled squares) and female (open circles) flies infected by the fungus *Entomophthora* over the season 2002.

**Table 1:**
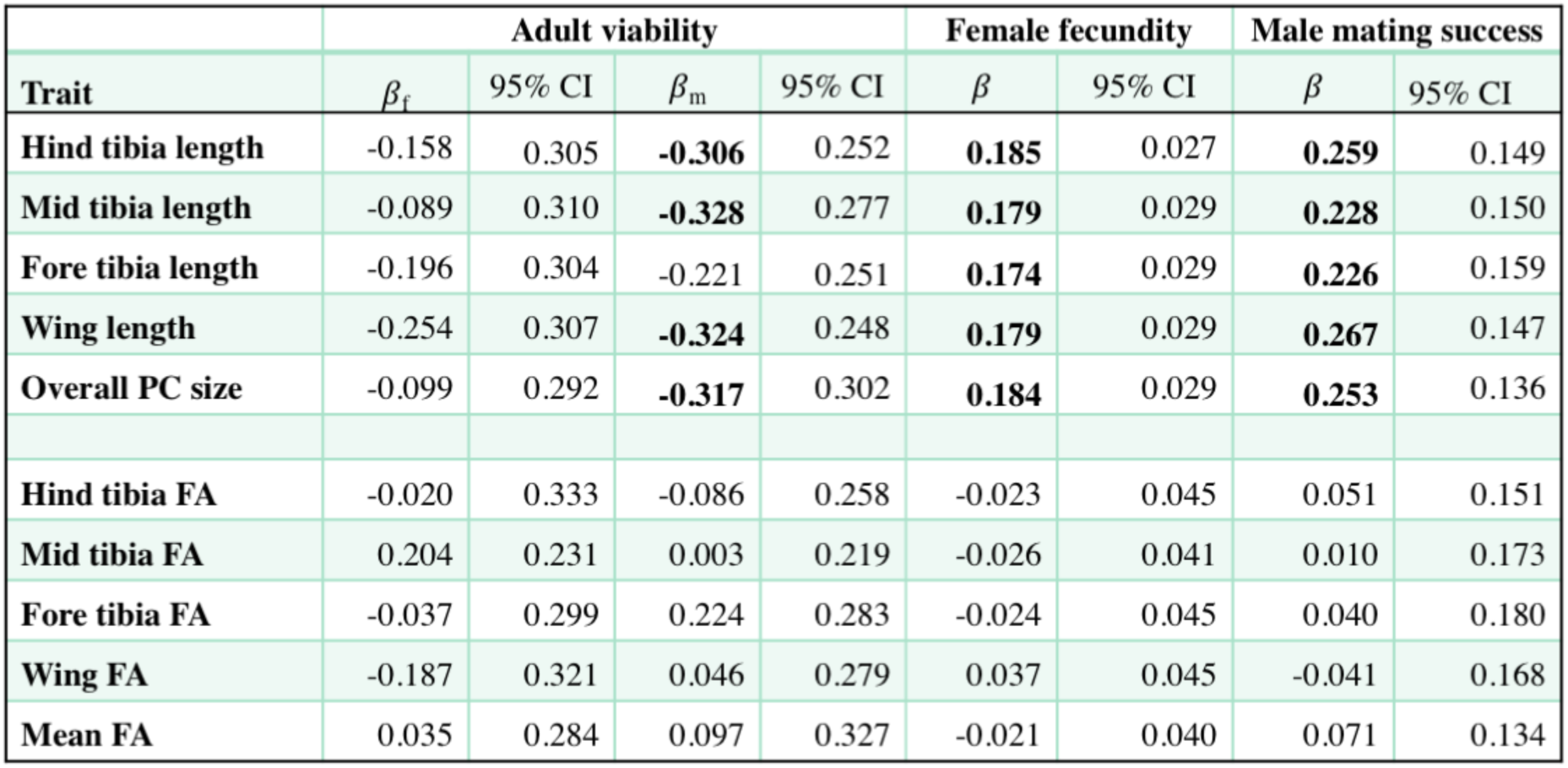
Overall intensities (*β* ± 95% CI) of female and male adult viability selection (N_f_ = 171 & N_m_ = 370) exerted by the fungus *Entomophthora*, female fecundity selection (clutch size; *N*_*f*_ = 126), and male sexual selection (pairing success; *N*_*m*_ = 370) for one Swiss population of yellowdung flies *(Scathophaga stercoraria)* over the season 2002. Significant coefficients are in bold (*P* < 0.001).

| Trait | Adult viability |  |  |  | Female fecundity |  | Male mating success |  |
| --- | --- | --- | --- | --- | --- | --- | --- | --- |
| | $\beta_f$ | 95% CI | $\beta_m$ | 95% CI | $\beta$ | 95% CI | $\beta$ | 95% CI |
| Hind tibia length | -0.158 | 0.305 | **-0.306** | 0.252 | **0.185** | 0.027 | **0.259** | 0.149 |
| Mid tibia length | -0.089 | 0.310 | **-0.328** | 0.277 | **0.179** | 0.029 | **0.228** | 0.150 |
| Fore tibia length | -0.196 | 0.304 | -0.221 | 0.251 | **0.174** | 0.029 | **0.226** | 0.159 |
| Wing length | -0.254 | 0.307 | **-0.324** | 0.248 | **0.179** | 0.029 | **0.267** | 0.147 |
| Overall PC size | -0.099 | 0.292 | **-0.317** | 0.302 | **0.184** | 0.029 | **0.253** | 0.136 |
| Hind tibia FA | -0.020 | 0.333 | -0.086 | 0.258 | -0.023 | 0.045 | 0.051 | 0.151 |
| Mid tibia FA | 0.204 | 0.231 | 0.003 | 0.219 | -0.026 | 0.041 | 0.010 | 0.173 |
| Fore tibia FA | -0.037 | 0.299 | 0.224 | 0.283 | -0.024 | 0.045 | 0.040 | 0.180 |
| Wing FA | -0.187 | 0.321 | 0.046 | 0.279 | 0.037 | 0.045 | -0.041 | 0.168 |
| Mean FA | 0.035 | 0.284 | 0.097 | 0.327 | -0.021 | 0.040 | 0.071 | 0.134 |

Fecundity selection (based on clutch size) on female body size was significantly positive, as is typical in this species (Jann et al., 2000; Kraushaar & Blanckenhorn, 2002). The intensity of fecundity selection, i.e. the slope relating relative clutch size to standardized body size (PC), varied significantly but unsystematically over the season (Table 1; interaction test: *F*_*6,109*_ = 3.50, *P* = 0.03). These estimates refer only to uninfected flies because all females infected with the fungus died before laying eggs, and therefore do not estimate fecundity selection exerted by the fungus beyond the parasite’s effect on adult mortality.

As usual in yellow dung flies, larger males had a mating advantage (Jann et al., 2000; Kraushaar & Blanckenhorn, 2002; Blanckenhorn et al., 2003; main effect of body size (PC): ***X*^2^** = 14.23; *P* < 0.001), while mean FA of all paired appendages did not affect male mating success(***X*^2^**= 1.17; *P* = 0.279; Table1; Fig. 2). Except for one seasonal sample on 29 May, the large male advantage was consistent throughout the season such that sexual selection intensity did not significantly vary across the season (body size by season interaction: ***X*^2^** = 0.11; *P* = 0.920; Table 1; Fig. 2). Interestingly, this pattern of positive sexual selection was nullified by fungal infection in those 28 (of a total of 47) infected males (of a total of 370 males, of which 148 had a mate) found mating in the field that later succumbed to the fungus (main effect of fungal infection: ***X*^2^** = 7.61; *P* = 0.006; fungus by size interaction: ***X*^2^** = 3.75; *P* = 0.053; Fig. 3).

**Figure 2:**
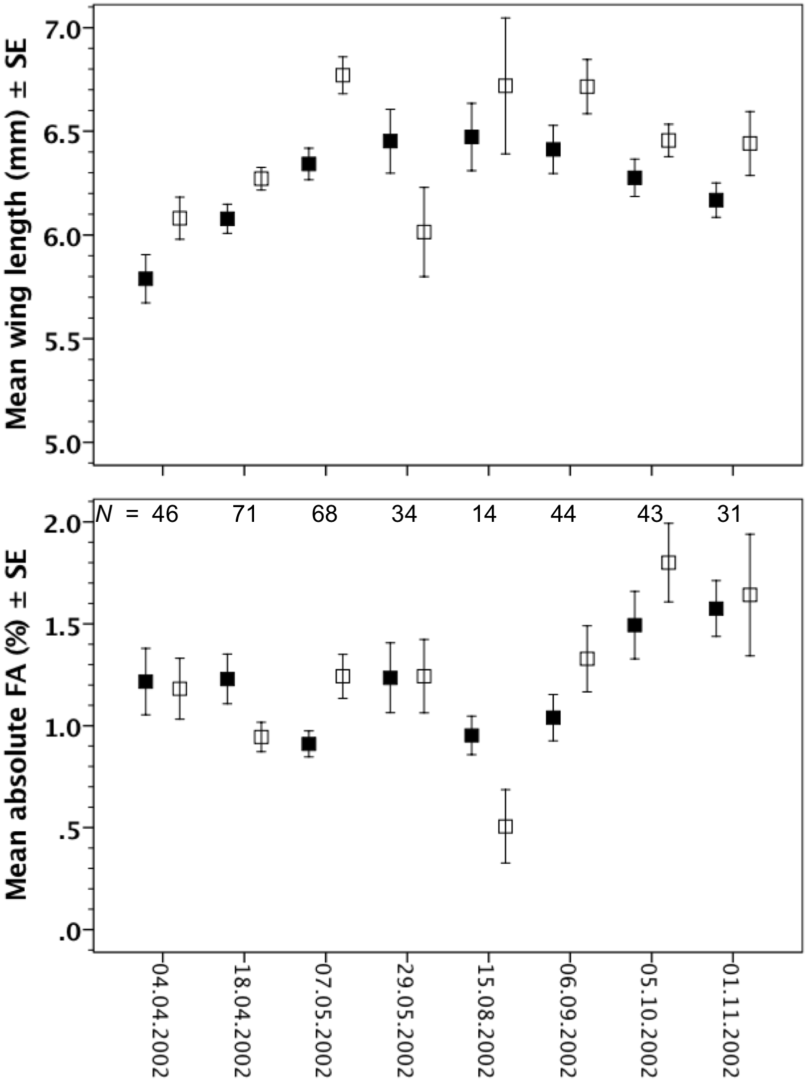
Body size (top; here exemplified by wing length) and mean percentage of fluctuating asymmetry (FA; bottom) of all traits for unpaired (filled squares) and paired males (open squares) over the season.

Table 1 gives directional (linear) selection coefficients, *β*, for the overall sample. Nonlinear (quadratic) selection coefficients, *γ*, were mostly low and not significant and are therefore not presented in Table 1. The only exception was female fecundity selection on body size, for which *γ* = 0.056 ± 0.025 was significantly positive, signifying accelerating selection (which has been reported before: Blanckenhorn, 2007, 2009). Leg and wing lengths were expectedly highly correlated in both sexes (range of bivariate correlations: *r* = 0.887 to 0.972), whereas FA of the legs and wings were largely uncorrelated (range: *r* = −0.024 to +0.215). Measurement of all paired traits was generally repeatable using our methods (*R* = 0.83 - 0.97), which was also true for asymmetry (R = 0.53 - 0.61), so that FA could be discerned from measurement error (all side by individual interactions *P* <0.01), fulfilling the criteria of proper FA assessment (Palmer & Strobeck, 1986; Knierim et al., 2007).

## Discussion

At our Swiss study population, the entomophagous fungal parasite *Entomophthora scatophagae*, which has been described as a specific parasite of adult yellow dung flies at several sites in Europe and North America (Hammer, 1941; Steinkraus & Kramer, 1988; Maitland, 1994; Steenberg et al., 2001), shows high and generally fatal infection rates of up to 50% during the cooler and more humid periods of the year. I here documented that this fungus exerts relatively strong and consistent negative viability selection on female and male adult body size in *S. stercoraria* in those parts of the year, when dung flies are most common (Table 1). Fungal infection further nullifies the usual large male mating advantage in sexual selection (Borgia, 1982; Jann *et al.* 2000; Blanckenhorn et al., 2003; Fig. 3), but does not affect female fecundity beyond its impact on mortality. This represents the first evidence demonstrating viability disadvantages of large yellow dung flies mediated by a parasite, which is generally rare in animals and particularly invertebrates (Blanckenhorn, 2000; Kingsolver & Pfennig, 2004; Gotanda et al., 2015). These results complement previous evidence of viability disadvantages of large flies at the juvenile stage, and lend further credence to the notion that the male-biased sexual size dimorphism of yellow dung flies is indeed roughly at evolutionary equilibrium (Blanckenhorn, 2007).

**Figure 3:**
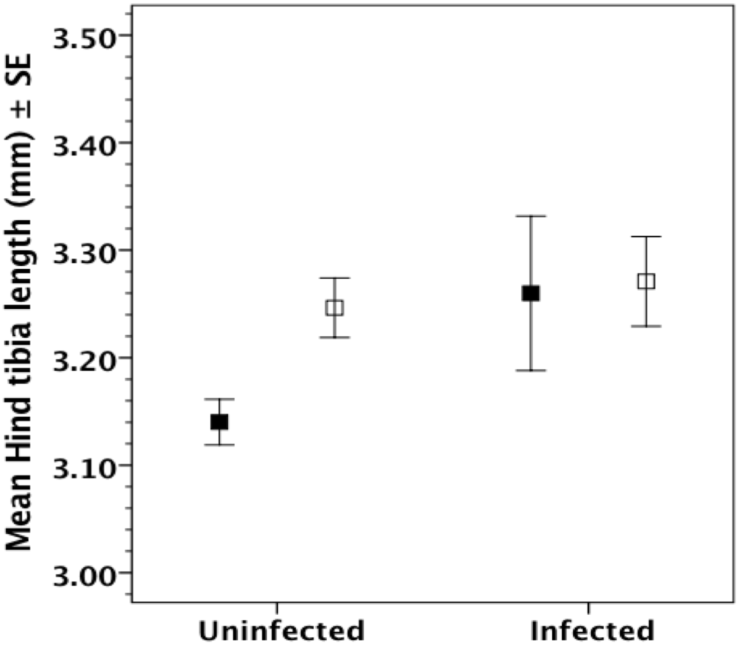
Body size (here exemplified by hind tibia length) of unpaired (filled squares) and paired males (open squares) when they were infected by the fungus or not (all seasonal samples combined).

This study is merely phenomenological, meaning that mechanisms were not assessed. Nonetheless, I speculate that the reduced parasite resistance of large flies signifies a trade-off between body size and immunity (Rantala & Roff, 2005; Schwarzenbach & Ward, 2006; Cotter et al., 2008). Based on age grading by wing injuries, Burkhard et al. (2002) found that adult age (i.e. longevity) of yellow dung flies in the field tends to be positively related to body size. Energy reserves also scale positively with body size (Reim et al., 2006; Blanckenhorn et al., 2007) and positively influence mating success (Blanckenhorn et al., 2003). Adult longevity under most environmental circumstances, including complete starvation, can therefore generally be expected to increase with body size on physiological grounds. One possible mechanism selecting against large body size is size-selective predation and/or parasitism (Blanckenhorn, 2000). However, beyond expectations of a rough positive correlation between predator and prey size (Brose et al., 2006; Vucic-Pestic et al.,2010), evidence for systematic size-selectivity of predators is generally weak at best, also for yellow dung flies (Blanckenhorn, 2000; Blanckenhorn et al., unpublished data). Size-selective parasitism has been reported in some parasitoids (McGregor & Roitberg, 2000), but otherwise few data exist (Zuk & Kolluru,1998; Blanckenhorn, 2000). Rather than invoking selective attraction of the parasite by larger hosts, I suspect that the ability to combat the parasite is compromised in larger flies due to their generally greater absolute energy demands in a stressful environment (trade-off hypothesis: Rantala & Roff, 2005; Reim et al., 2006; Schwarzenbach & Ward, 2006, 2007; Cotter et al., 2008). Females were infected more by the parasite here (Fig. 1), probably related to their generally greater reproductive burden (i.e. the cost of producing expensive eggs rather than cheap sperm; cf. Nunn et al., 2009; but see e.g. Rantala et al., 2007, for opposite results). Nevertheless, the standard sex differences in reproductive (energetic) costs should be somewhat offset by the male-biased sexual size dimorphism of yellow dung flies, which implies relatively greater costs when producing and maintaining the larger and condition-dependent male body (Blanckenhorn, 2000, 2007), and might explain why size-dependent viability selection here turned out to be steeper in males (Table 1). Yellow dung fly females indeed produce higher heritable levels than males of phenoloxidase (PO; Schwarzenbach et al., 2005), one of the central mediators of insect immunity (Schmid-Hempel, 2005; Rolff & Reynolds, 2009; González-Santoyo & CÓrdobaAguilar, 2012), and higher PO levels decrease adult longevity in this species, demonstrating a trade-off (Schwarzenbach & Ward, 2006). However, higher PO levels did not lead to greater resistance against mites or another fungus (Schwarzenbach & Ward, 2007), and PO is also unrelated to body size in *S. stercoraria* (Schwarzenbach et al., 2005). Overall, therefore, the evidence in favor of immunity mediating the higher mortality of large-bodied dung flies here remains limited.

In contrast to body size, fluctuating asymmetry (FA) of legs and wings influenced none of the fitness components investigated here (contrary to Liggett et al., 1993, but confirming Blanckenhorn et al’s, 2003, earlier sexual selection study). I had expected, based on evidence in other animals (Rantala et al., 2000, 2004), that low FA would be a signal of greater immunocompetence augmenting resistance against parasites, but this was not found. It is not unlikely that FA is a bad indicator of developmental stability in general, as various reviews have revealed no clear verdict based on the available evidence on this question, so the entire concept remains controversial (Møller & Thornhill, 1997; Møller & Swaddle, 1997; Palmer, 2000; various articles in Polak, 2003; VanDongen, 2006; Knierim et al., 2007). In yellow dung flies, beyond Liggett et al. (1993) the evidence for a role of FA in sexual selection is close to nil (Blanckenhorn et al., 2003; Blanckenhorn & Hosken, 2003; this study). What remains is that FA reliably indicates at least hot temperature stress in this species (Hosken et al., 2000; Blanckenhorn & Hosken, further unpublished data), even though Floate & Coughlin (2010) concluded that FA is no good biomarker of toxic chemical residues in cattle dung. As the survival of flies infected with the fierce entomophagous fungus *Entomophthora scatophagae* here was related to the size (discussed above) but not the fluctuating asymmetry of wings and legs, I conclude that FA is no good indicator of immunocompetence in yellow dung flies either (Rantala et al., 2000, 2004, 2007; Rantala & Roff, 2005; Yourth et al., 2002; Schwarzenbach & Ward, 2006; Cotter et al., 2008).

## Acknowledgements

I thank the University of Zurich and the Swiss National Fund for financial support over the years, and Ana, Lukas, Nils and Patricia for doing the measurements.

